# Transcriptional and metabolic stasis define desiccation-induced dormancy in the soil bacterium *Arthrobacter* sp. AZCC_0090 until water vapor initiates resuscitation

**DOI:** 10.1101/2025.10.16.681527

**Authors:** Paul Carini, Adriana Gomez-Buckley, Christina R. Guerrero, Melanie R. Kridler, Isabella A. Viney, Roya AminiTabrizi, Malak M. Tfaily, Peter Moma, Laura K. Meredith, Katherine B. Louie, Benjamin P. Bowen, Trent R. Northen, Oona Snoeyenbos-West, Ryan P. Bartelme

**Author notes:** Authors contributed equally to this work (Adriana Gomez-Buckley, Christina R. Guerrero, Melanie R. Kridler, Isabella A. Viney).

## Abstract

Microbes inhabiting soils experience periodic water deprivation. The effects of desiccation on DNA, protein, and membrane integrity are well-described. However, the effects of drying and rehydration on the composition of cellular RNA and metabolites are still poorly understood. Here, we describe how slow drying and rehydration with water vapor influence the composition of RNAs and metabolites in a soil *Arthrobacter*. While drying reduced cultivability relative to hydrated controls, water vapor rehydration fully restored it. Ribosomal RNA proportions remained constant throughout all treatments, and mRNA profiles showed stable composition during desiccation—changing only during transitions into and out of desiccation-induced dormancy. Six transcriptional modules displayed distinct expression patterns in desiccated-rehydrated samples relative to hydrated controls, including desiccation-rehydration responsive and rehydration-specific profiles. Targeted intracellular metabolomics revealed similarly static profiles during desiccation, with a cluster of ribonucleosides and nucleobases increasing in response to desiccation and returning to baseline levels upon rehydration with water vapor.

These findings demonstrate that both mRNA and metabolite profiles remain essentially frozen in desiccated *Arthrobacter*, with dynamic changes occurring only during state transitions. These results have important implications in environments with frequent drying cycles where stable mRNA in dormant cells combined with intracellular RNA recycling may obscure interpretations of RNA-based environmental analyses that use RNA as a marker of microbial activity. Our results suggest that RNA-based activity assessments in periodically dry environments require careful consideration of dormancy-associated molecular preservation.

**SIGNIFICANCE STATEMENT:** Metabolic activity quickly ceases in drying bacteria as they enter desiccation-induced dormancy. We show mRNA and metabolite profiles were variable during drying and rewetting but did not change while desiccated. Additionally, water vapor stimulated the shift from the static to active state when exiting desiccation-induced dormancy. These shifts coincided with increased cultivability, indicating water vapor resuscitated dry cells. Because RNAs are transient, labile molecules that are turned over rapidly in growing bacteria, the presence of RNA in the environment is used as a marker for microbial activity. Our research shows this assumption may not hold for desiccated cells, indicating reliance on RNA as a marker of activity in environments that experience drying may obscure estimates of *in situ* microbial activity.

## INTRODUCTION

Water is required for life as a substrate in essential biological reactions and as a solvent to transport nutrients and waste (1). As such, the lack of water stresses nearly all organisms. Future climate models predict changes in global precipitation patterns, further restricting precipitation in drylands and introducing drought to regions that have not previously experienced it (2–4). The impacts of drought can accelerate agricultural crop losses and food insecurity (5, 6), reduce freshwater connectivity and services (7), and alter soil biogeochemistry (8–11). Soil microbiomes are restructured during drought (8, 12, 13) and can influence crop drought tolerance (14–17). Despite the impacts of drying on soil microbes and their critical roles in global carbon dynamics and food production, basic aspects of how soil microbes persist in the dry state remain unknown.

As soil dries, the solutes in pore water become concentrated, reducing water potential and limiting water availability. Microbes equilibrate rapidly to the water potential of their surroundings because of their small size (13, 18). Eventually, growth substrates precipitate, rendering them unavailable for microbial metabolism. Thus, microbial activity slows with dehydration until cellular respiration stops (13, 19). Macromolecules in dehydrated cells are prone to structural damage. Proteins fold improperly, denature, or are oxidized (20–22). DNA can also be oxidized by reactive oxygen species (ROS) (18, 23). And membrane integrity can be compromised in the desiccated state or upon rehydration, where lysis may occur due to rapid turgor pressure changes that come with the influx of water (24, 25).

Some bacteria endure water scarcity through dormancy, a reversible state of reduced metabolic activity (26). Spore-forming microbes like *Bacillus* and *Streptomyces* undergo dormancy via cell differentiation to produce ultra-resistant spores in response to nutrient starvation (26–28). Yet, many—perhaps most—soil microbes cannot sporulate. Instead, these non-spore formers employ anhydrobiosis, a state of suspended animation where organisms survive extreme water loss by halting metabolism to nearly undetectable levels while maintaining cellular integrity (29, 30).

Anhydrobiosis enables survival for years to decades in the dry state with rapid metabolic resumption upon rehydration. While well characterized in tardigrades, nematodes, and some plants, the molecular mechanisms underlying anhydrobiosis in non-spore forming soil microbes remain less understood (29–31). Non-spore formers respond to water loss by mitigating cellular damage through multiple coordinated mechanisms. These include accumulating organic osmolytes like trehalose that lower intracellular solute potential and may stabilize membranes and macromolecules (13, 32–36); remodeling membrane lipids to resist disruption (37, 38); producing exopolysaccharides and biofilms that slow water loss (39, 40); upregulating oxidative stress protection (31, 41, 42); and enriching transcripts for molecular chaperones and DNA repair proteins (41–43).

Yet, the fate of RNA and intracellular metabolites during desiccation-induced dormancy remain understudied, particularly how the transcriptional and metabolic profiles change in dry cells and upon rehydration. While endospores show a dramatic reduction in RNA content during dormancy (44), the RNA dynamics in desiccation-tolerant non-spore-forming bacteria are underexplored. This question is important to resolve for environmental microbiology, where RNA abundances—both ribosomal and messenger RNA—serve as a key indicator of microbial activity across diverse environments (45–48), including soils (49–51). If RNAs are detectable in inactive cells, RNA may not reliably indicate microbial activity. Here, we demonstrate that desiccation induces a dormant state in a soil *Arthrobacter* characterized by: 1) reduced but recoverable cultivability, with water vapor alone sufficient for resuscitation; 2) stable mRNA and intracellular metabolite profiles during desiccation, despite dynamic changes during transitions into and out of the desiccated state; and 3) ribosomal RNAs that occupy a constant fraction of the total RNA pool irrespective of hydration (and hence, activity) status. By identifying distinct temporal patterns in mRNAs and metabolites across the desiccation-rehydration cycle, we reveal implications for how RNA-based approaches may misrepresent microbial activity in environments that experience periodic drying.

## RESULTS & DISCUSSION

### Experimental design

To investigate cellular responses to desiccation stress, we studied *Arthrobacter* sp. strain AZCC_0090, a soil actinobacterium isolated from semiarid soil in Southern Arizona that is closely related to the *Arthrobacter* phylotypes found across the United States (52–54). We developed a controlled desiccation system using two humidity chambers in which the relative humidity (RH) of the atmosphere was controlled with saturated salts or ultrapure (Nanopure) water (Fig. 1 a,b). The chambers consisted of a “control” chamber maintained at 100% RH and a “treatment” chamber where RH was gradually reduced from 100% to 26% over 14 days to desiccate cells, corresponding to an atmospheric water potential change from 0 to -183 MPa (Fig. 1c). After 14 days of desiccation, we restored the 100% RH atmosphere of the treatment chamber to rehydrate dried cells for two additional days (Fig. 1c).

**Figure 1:**
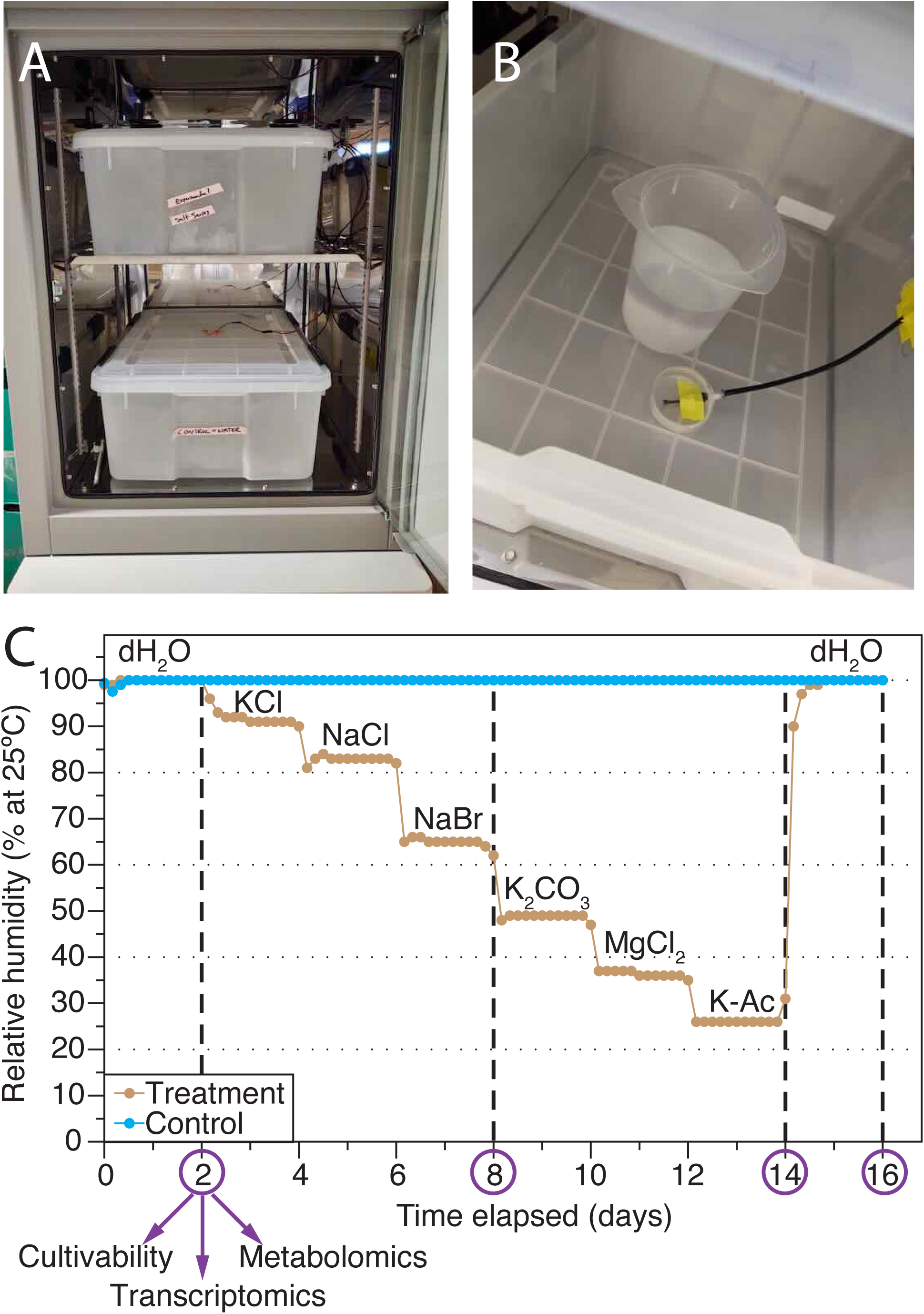
Experimental system for controlled desiccation and rehydration. (A) Custom humidity chambers housed within a temperature-controlled incubator. (B) Interior chamber configuration showing saturated salt solution for humidity control and real-time temperature/humidity sensor. (C) Relative humidity profiles over the 16-day experiment, with sampling points (purple circles and vertical dashed lines) and water/salt solutions used at each stage. Abbreviations: K-Ac: Potassium Acetate; dH2O: deionized water.

The degree of desiccation extended far below the water potentials that inhibit growth in both *E. coli* (-4.6 MPa=96.7% RH) and *Arthrobacter spp*. (-17 MPa=88.0% RH) (18). Cells were collected on non-hygroscopic polycarbonate filters and placed into these chambers. Replicate filters were collected from the chambers at four timepoints to analyze cell cultivability, intracellular metabolite profiles, and gene expression (Fig. 1c).

This experimental design incorporated three unique aspects relative to previous studies (41, 42). First, hydrated control cells were maintained alongside treatments, allowing us to specifically identify the responses to desiccation and rehydration. Second, dehydration occurred gradually over 14 days rather than rapidly. Finally, rehydration was achieved through water vapor alone. This approach enabled control over cellular water status while minimizing confounding variables as suggested by Bosch, et al (29).

### Dry cells are dormant

We defined cultivability as the fraction of filtered cells that were culturable on solid media. Initial cultivability immediately after filtering was 36.3 ± 24.5% (mean ± SD, n=4; Fig. 2A), with variability reflecting the technical challenges of quantitatively recovering cells from filters. Cells maintained in hydrated control conditions showed stable cultivability throughout the experiment, with no significant variation over time (Kruskal Wallis p = 0.580; Fig. 2A). In contrast, cells subjected to desiccation showed significant temporal variation in cultivability (Kruskal Wallis p = 0.022), with mean values decreasing by an order of magnitude from 24.0% at day 2 (100% RH) to 2.46% at day 8 (65% RH; Fig. 2A). Desiccated cells showed significantly reduced cultivability compared to hydrated controls at both day 8 and day 14 (Mann-Whitney p = 0.028 for both timepoints; Fig. 2A), indicating that reduced cultivability was due to desiccation.

**Figure 2:**
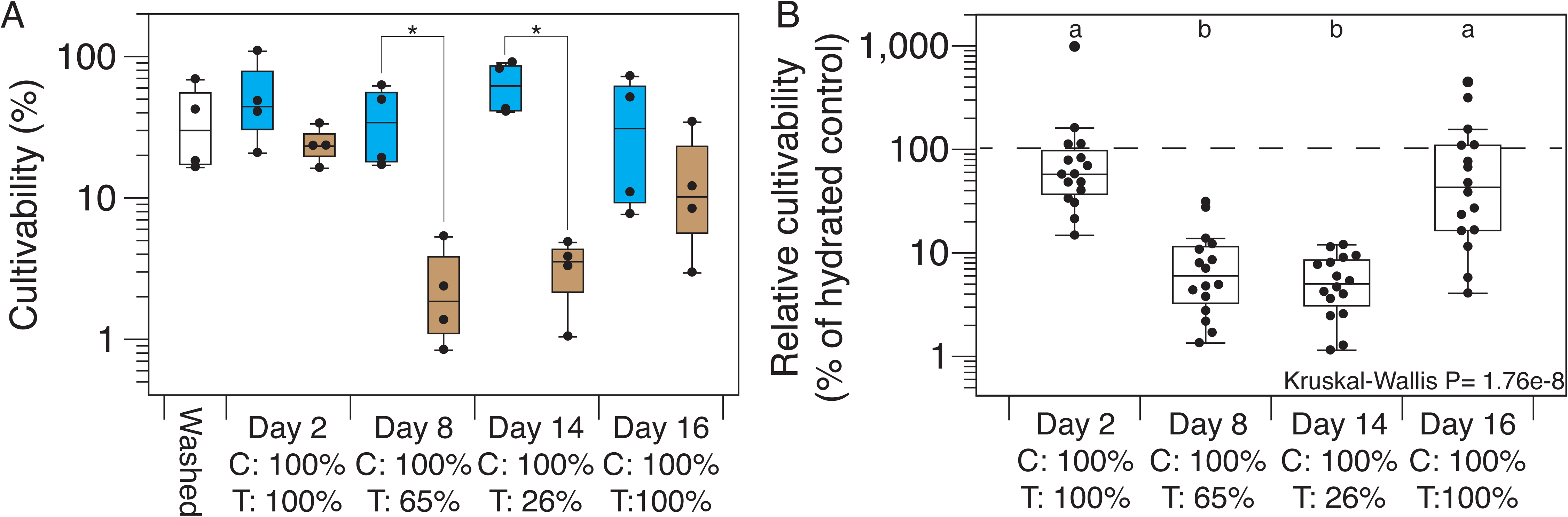
Desiccation-induced growth inhibition is reversible with water vapor. (A) Cell cultivability throughout the 16-day experiment with points showing individual replicate filters (n=4) for washed cells (filtered and assayed immediately without incubation; no fill), controls (blue), and treatment samples (brown). Brackets with asterisk indicate significant differences between treatment and control (Mann-Whitney p ≤ 0.05). (B) Relative cultivability (treatment/control ratio) showing desiccation-induced decreases in cultivability and recovery post rehydration. Points are all-by-all pairwise cultivability ratios. The dashed line in (B) indicates 100% relative cultivability (treatment equals control). Box plots with shared letters are not significantly different (Dunn’s test with Bonferroni correction, p > 0.05). C=Control, T=Treatment, given in terms of percent RH.

Water vapor restored cultivability of desiccated cells, demonstrating dry cells were dormant, not dead. When cells from day 14 (26% RH) were exposed to 100% RH for two days, their mean cultivability increased to 14.4%—statistically indistinguishable from constantly hydrated controls at day 16 (Mann-Whitney P = 0.49; Fig. 2A). Although the AZCC_0090 genome lacks genes associated with sporulation (53), we tested whether the observed recovery was due to spore formation. No growth was observed after treating cells with ethanol, confirming AZCC_0090 survives desiccation without sporulating.

We further calculated the relative effects of desiccation and rehydration as the cultivability ratio of treatment to control cells at each timepoint (Fig. 2B). This confirmed significant variation across the experiment (Kruskal-Wallis p = 1.76e-8), with days 8 and 14 showing significantly lower relative cultivability compared to both pre-desiccation (day 2) and post-rehydration (day 16) timepoints (Fig. 2B; Dunn’s test, Bonferroni-adjusted P ≤ 0.05). These results demonstrate that desiccation caused reduced cultivability and water vapor restored cultivability in desiccated *Arthrobacter*.

### Effect of desiccation and rehydration on RNA content and composition

Initial total RNA yields from washed cells were 150 ± 29.1 ng µl^-^¹ (mean ± SD, n=3; Fig 3a). After 2 days at 100% RH, total RNA decreased in both the control and treatment groups relative to the washed cells, with no significant difference between the conditions, likely due to nutrient starvation leading to ribosome degradation (55, 56). Subsequently, RNA amounts in the control condition varied significantly over time (Kruskal-Wallis p = 0.033 between days 2 and 16; Fig. 3A), while desiccated and rehydrated cells showed no temporal variation (p = 0.305; Fig. 3A). Because rRNA constitutes most of the total RNA, these temporal differences likely represent ribosomal turnover in metabolically active control cells versus desiccated cells undergoing metabolic arrest. However, no significant differences in total RNA content were detected between control and treatment groups at any individual timepoint (Wilcoxon rank-sum tests, all p ≥ 0.08).

**Figure 3:**
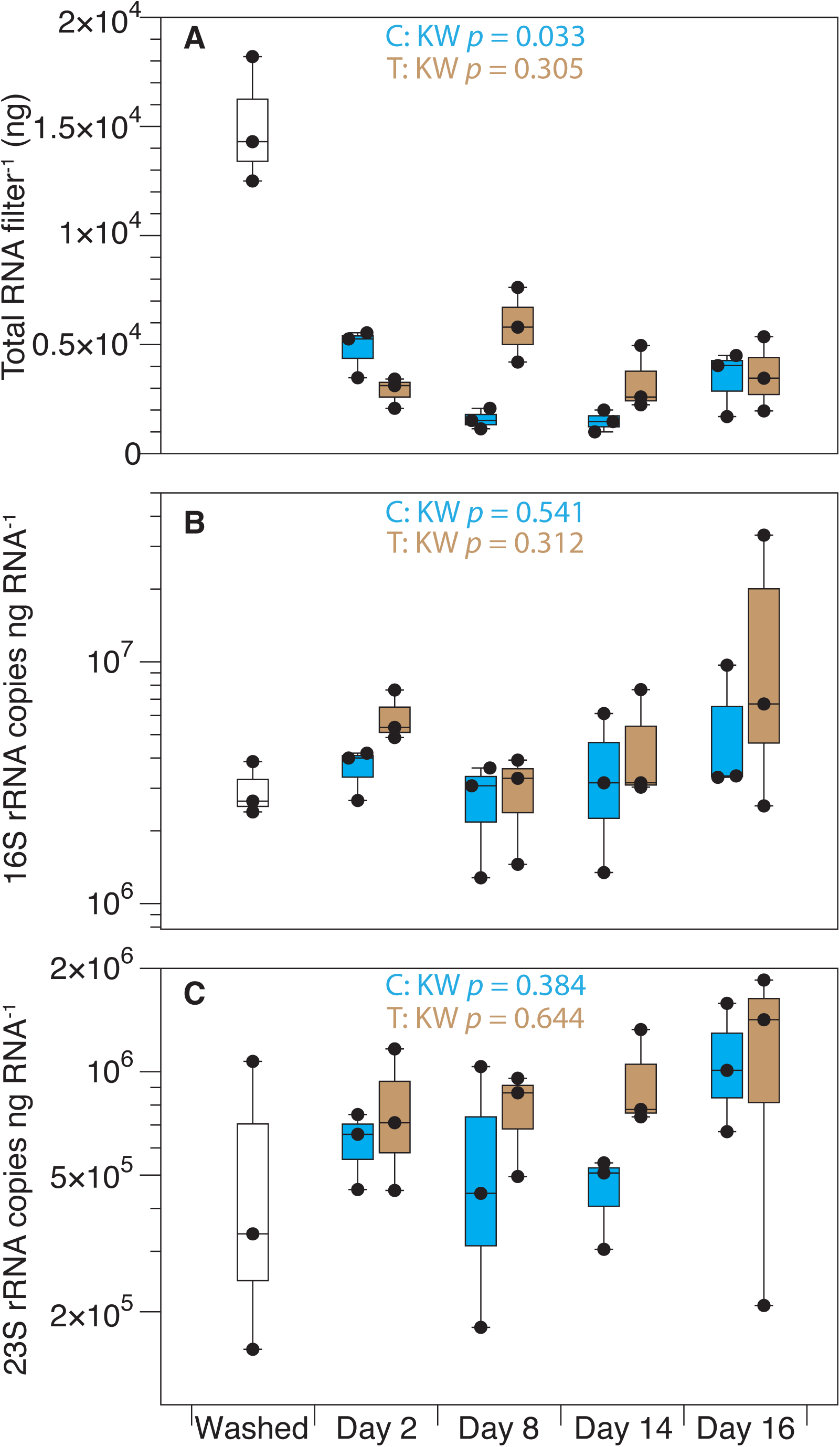
rRNA composition during desiccation and rehydration is constant. (A) Total RNA yield from filtered cells over the 16-day experiment. Individual points represent RNA concentrations from replicate filters: washed cells (filtered and assayed immediately without incubation; no fill), hydrated controls (blue), and desiccation-rehydration-treated samples (brown). (B,C) Normalized abundance of rRNA transcripts (copies ng total RNA ¹) for (B) 16S rRNA and (C) 23S rRNA. Box plots show median (horizontal line), interquartile range (box), and 1.5 × interquartile range (whiskers). Kruskal-Wallis test p-values shown for control (C) and treatment (T) groups across all timepoints (Days 2-16).

We investigated whether the fraction of total RNA occupied by the main cellular rRNAs (16S and 23S rRNA) varied during the experiment. To do this, we quantified 16S and 23S rRNA transcript abundances and normalized them to the amount of total extracted RNA (Fig. 3 B,C). The proportions of both rRNA species remained stable across all time points in both treatment and control conditions (Kruskal-Wallis 16S rRNA treatment *p*=0.312 control *p*=0.541; 23S rRNA treatment *p*=0.644, control *p*=0.384). Pairwise comparisons revealed no significant differences between control and treatment groups at any individual timepoint for either 16S rRNA or 23S rRNA (Wilcoxon rank-sum tests, p ≥ 0.1). These results indicate that the contributions of 16S and 23S rRNAs to the total RNA pool were maintained despite changes in total RNA content.

In contrast to the effects on rRNA, drying and rehydrating reshaped mRNA transcription patterns, but transcription profiles were stable in desiccated cells. Analysis of mRNA transcriptional responses revealed significant effects of both time point and treatment, with each factor explaining ∼20% of the observed variation in gene expression (PERMANOVA time: 21.9%, *p*=0.001; treatment: 19.1%, *p*≤0.001; Supplemental Fig. 1). The interaction between time and treatment explained an additional 16.8% variation (*p*=0.009), indicating that temporal gene expression patterns differed between control and treatment conditions. The control samples showed changes over time, though none of the consecutive timepoint shifts were significant (PERMANOVA p >0.05 between consecutive timepoints). In contrast, the treatment samples displayed a significant shift in gene expression upon desiccation (day 2 to 8: PERMANOVA p=0.023, R²=0.53). Upon rehydration, although the time variable explained a large proportion of variance in gene expression profiles (R²=0.56), this shift was not significant (day 14 to 16: PERMANOVA p=0.1). Yet, the transcriptional profiles were indistinguishable in the desiccated state (day 8 to 14 PERMANOVA p=0.73, R^2^= 0.15). These findings show the composition of mRNA profiles were variable during drying and rehydration but did not change over time in dry *Arthrobacter* cells.

Analysis of the temporal patterns in gene expression supported our interpretation that gene expression profiles remained static during desiccation while hydrated cells were transcriptionally dynamic. We detected 1,409 genes with significantly different temporal expression patterns across treatment and control conditions (ImpulseDE2 FDR-corrected p ≤0.05; Supplementary Table 1). These genes clustered into six distinct gene co-expression modules based on their combined temporal patterns across both conditions (Fig. 4, Supplementary Table 1). Across all transcriptional modules, we observed further evidence of a frozen transcriptional state in desiccated cells, as no substantial change in mean gene expression patterns in the treatment samples between days 8 and 14 was apparent (bold brown lines, Fig. 4). However, transcriptional profiles were dynamic in treatment samples during drying and/or rehydration in Transcriptome Modules 2, 3, 4, 5 and 6 (Fig. 4 B-F). In contrast, these gene co-expression modules exhibited distinct variable responses throughout the experiment in control samples, including between days 8 and 14, suggesting hydrated cells were transcriptionally active and dynamically responded to starvation.

**Figure 4:**
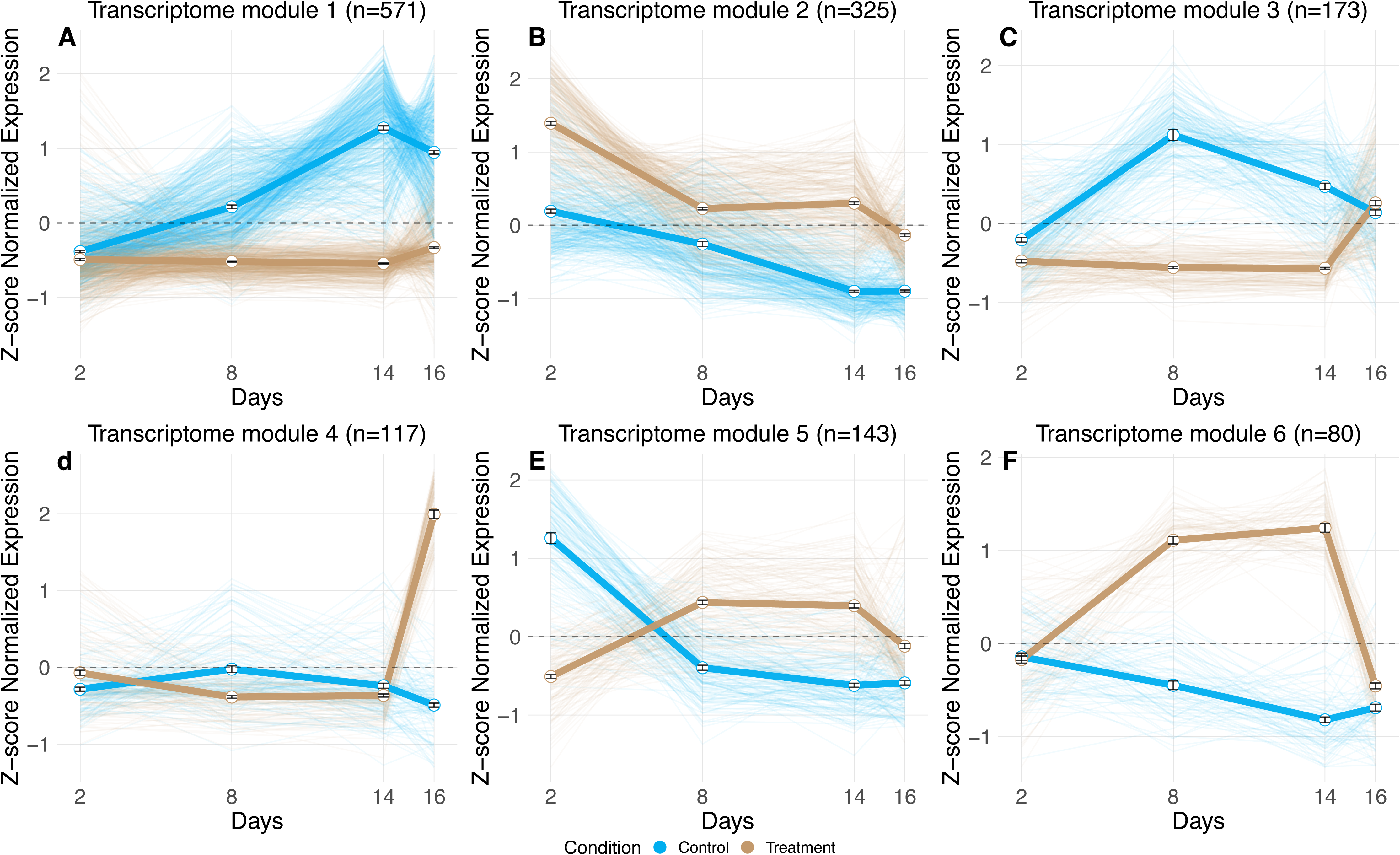
Temporal expression modules of differentially expressed genes. Six distinct co-expression modules (A-F) identified through correlation-based hierarchical clustering of gene transcripts with significantly distinct temporal responses across treatment and control conditions (ImpulseDE2, FDR-corrected p ≤ 0.05). Each panel displays mean z-score normalized gene expression value (±SEM) for control (bold blue) and treatment (bold brown) conditions, with individual gene profiles shown as thin lines. Sample sizes (n) indicate the number of expressed genes within each module.

Two main desiccation/rehydration-specific gene expression patterns emerged in the treatment samples. First, Transcriptome Modules 5 (143 genes) and 6 (80 genes) exhibited similar temporal responses where expression increased during drying, remained elevated throughout desiccation, and returned to baseline after rehydration (Fig. 4 E,F and Supplementary Table 1). While these transcriptional modules showed similar expression patterns in treatment samples, their expression in hydrated controls was distinct, suggesting that although their roles are similar during desiccation, they were expressed independently during starvation.

The annotations of genes expressed in Transcriptome Modules 5 and 6 revealed responses consistent with desiccation tolerance mechanisms in non-spore forming microbes (Fig. 4 E,F; Supplementary Table 1). For example, Transcriptome Module 5 included genes coding for fatty acid metabolism, the osmolyte trimethyl glycine (betaine) synthesis and transport, and purine catabolism. The coordinated upregulation of choline-to-betaine conversion machinery alongside osmolyte transporters suggests AZCC_0090 depends on the conversion of external choline to betaine for osmoregulation during drying. The upregulation of compatible solute synthesis and transport machinery is a common response to drying (31). Additionally, Transcriptome Module 5 contained regulatory and repair systems including numerous transcriptional regulators, DNA repair proteins, and ROS protection enzymes—all hallmarks of desiccation stress responses in microbes (31). Finally, a large-conductance mechanosensitive channel was also identified that may prime cells to cope with the stresses associated with rehydration (57). However, we were surprised by the presence of genes in Transcriptome Module 5 that code for protein synthesis machinery including several ribosomal proteins and translation factors. This was unexpected given the apparent metabolic slowdown during drying, though similar findings have been previously reported (36, 58). Transcriptome Module 6 focused on distinct cellular processes including carbohydrate processing, protein quality control through proteases, aromatic compound and lipid metabolism, and nucleotide processing via a putative NUDIX family pyrophosphohydrolase. The genes in Transcriptome Modules 5 and 6 largely align with previously reported bacterial desiccation responses, including expected mechanisms for ROS mitigation, osmoregulation, putative membrane remodeling, and activation of DNA and protein repair systems to address oxidized biomolecules (31, 36, 59). Notably, 33% of Transcriptome Module 5 and 38% of Transcriptome Module 6 genes lacked meaningful annotations, suggesting novel mechanisms and genes may contribute to desiccation tolerance. The expression patterns of both modules—increased relative abundance during drying and decreased relative abundance upon rehydration—suggest these transcripts are likely recycled upon rewetting.

The second desiccation-rehydration specific gene expression pattern was the dramatic increased abundance of 290 genes across Transcriptome Modules 3 & 4 during rehydration with water vapor, suggesting key roles in cellular resuscitation (Fig. 4 C,D). These genes were expressed at low levels in the treatment condition prior to rehydration though their expression was distinct in controls (Fig. 4 C,D and Supplementary Table 1). Transcriptome Module 3 contained genes coding fatty acid beta-oxidation and aromatic compound degradation pathways, alongside abundant transcriptional and translational machinery indicating active protein synthesis during rehydration. Stress response signatures included multiple chaperones and catalase, suggesting mitigation of oxidative damage, while the high density of transcriptional regulators indicates extensive gene expression reprogramming during rehydration. Transcriptome Module 4 contains genes coding for osmoprotectant transport (distinct from Module 5), a xylose metabolism operon, histidine catabolism, DNA repair and replication machinery, and energy metabolism genes. The xylose pathway and associated sugar phosphate enzymes may shuttle 5-carbon sugars toward phosphoribosyl pyrophosphate (PRPP) for nucleic acid repair or synthesis. Histidine degradation may serve dual functions: chelating divalent cations during desiccation to reduce ROS generation, then providing carbon and nitrogen upon rehydration (60). InterProScan analysis revealed seven ’hypothetical proteins’ (HNP00_000350, HNP00_000836, HNP00_001555, HNP00_001771, HNP00_002967, HNP00_002968, and HNP00_003348) containing disorder domains with polyampholyte subdomains—signatures of eukaryotic anhydrins with chaperone-like roles during desiccation (61, 62). Two of these (HNP00_001555 and HNP00_001771) also contained HNH-nuclease-like domains, suggesting dual roles in nucleic acid metabolism and protein stability. Together, Transcriptome Modules 3 and 4 represent coordinated cellular machinery activated during rehydration: Module 3 mobilizes lipids for energy while reactivating transcription, translation, and stress management, while Module 4 manages osmotic stress, alternative carbon utilization, DNA repair, and resource acquisition essential for growth resumption.

### Desiccated cells are in metabolic stasis

To identify metabolic changes that correspond to survival during drying and rehydration we also conducted targeted and untargeted intracellular metabolite analysis alongside the transcriptomes. Targeted metabolite analysis revealed that treatment was the dominant factor shaping metabolic composition (PERMANOVA R² = 24.3%, p < 0.001), followed by temporal dynamics (PERMANOVA R² = 17.8%, p < 0.001) and their interaction (PERMANOVA R² = 15.9%, p = 0.002; Supplemental Fig. 2). Collectively, these factors explained 58% of total targeted metabolomic variance in the experiment. Like the results from the transcriptome analysis, we observed a treatment-specific response characterized by metabolic restructuring during drying (between days 2 and 8; PERMANOVA R² = 63.8%, p = 0.01) and rehydration (between days 14 and 16; PERMANOVA R² = 33.3%, p = 0.01), with no change in the dry state (between days 8 and 14; PERMANOVA R² = 7.3%, p = 0.71). In contrast, control samples displayed a muted temporal drift (Day 2-8: PERMANOVA R² = 26.6%, p = 0.01; Day 8-14: PERMANOVA R² = 21.5%, p = 0.05; Day 14-16: PERMANOVA R² = 10.8%, p = 0.43). These results demonstrate that like the observed compositional stasis in the dry state transcriptomes, *Arthrobacter* also undergo metabolic dormancy during desiccation with subsequent recovery toward profiles like the pre-desiccated state.

To identify metabolites with correlated temporal profiles, we clustered targeted metabolite profiles based on the shape of their normalized peak height variation over time. Of the 105 targeted metabolites analyzed, 46 (43%) showed significantly distinct temporal patterns across the treatment and control samples (modified ImpulseDE2 FDR-corrected p ≤0.05; Supplementary Table 2). Clustering of significantly different metabolites across the treatment and control conditions revealed four Targeted Metabolite Modules (Fig. 5; Supplementary Table 2). The largest Targeted Metabolite Module (module 1, 21 metabolites) showed a similar profile during desiccation to that observed for Transcriptome Modules 5 and 6 (Figs 4 E,F), characterized by increasing peak area during drying, an elevated level when dry, and a return to baseline after rehydration (Fig. 5A). Of these metabolites, 12 (57%) were either directly related to nucleic acid metabolism, including all ribonucleosides, purine and pyrimidine bases, or nucleotide degradation products. These nucleotide degradation products likely accumulate because transcription ceases in the transition to the dry state. We speculate that existing nucleotide monophosphates are progressively dephosphorylated, and their N-glycosidic bonds cleaved, releasing the ribonucleosides and nitrogenous bases we observed. This observation is consistent with internal mRNA recycling across the desiccation-rehydration continuum. Also notable were the presence of potential antioxidant compounds including B-vitamins (B_1_ & B_3_), L-gulonolactone (an intermediate in the ascorbic acid biosynthetic pathway), and N-acetyl L-glutamic acid (63–66). Finally, both L-methionine and methylthioadenosine (MTA) were elevated during desiccation. These metabolites may work as part of a methionine-methionine sulfoxide antioxidant cycling system where methionine serves as a ROS scavenger, with MTA indicating ongoing methionine recycling, SAM production, and/or polyamine synthesis during desiccation (67).

**Figure 5:**
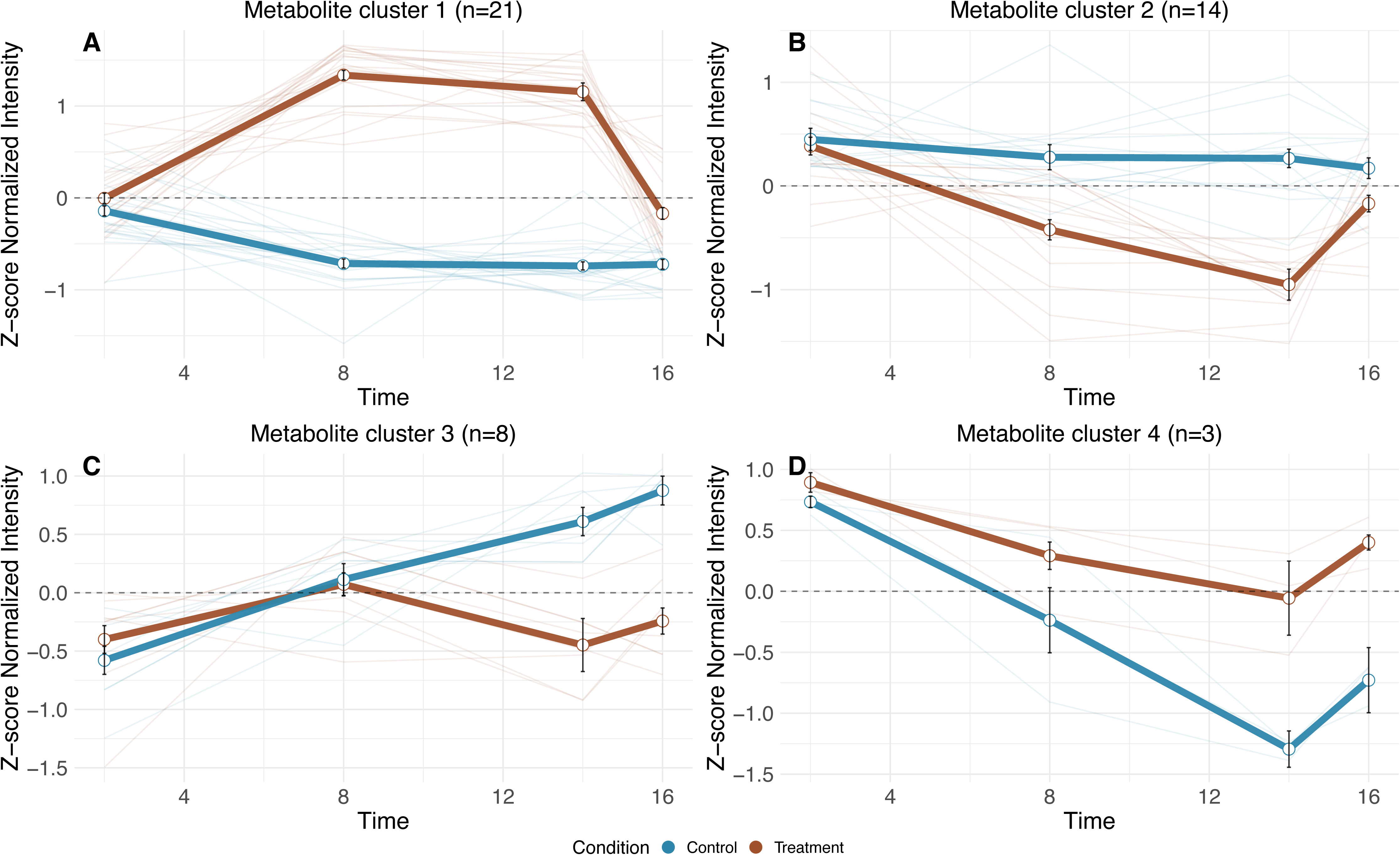
Temporal modules of differentially abundant targeted metabolites. Four metabolite modules (A-D) were identified through correlation-based hierarchical clustering of targeted metabolites with significantly distinct temporal responses across treatment and control conditions (modified ImpulseDE2, FDR-corrected p ≤ 0.05). Each panel displays mean z-score normalized targeted metabolite peak height value (±SEM) for control (bold blue) and treatment (bold brown) conditions, with individual metabolite profiles shown as thin lines. Sample sizes (n) indicate the number of targeted metabolites within each module.

The remaining Targeted Metabolite Modules showed metabolite depletion in treatment samples from days 2-14, potentially due to active catabolism, abiotic degradation, non-enzymatic transformations, or residual enzymatic activity (29)—possibilities our data cannot distinguish. Betaine, a known osmolyte, decreased in both conditions (Targeted Metabolite Module 4; Fig. 5D) but more steeply in controls. In desiccated cells, betaine synthesis transcripts were more abundant (Transcriptome Module 5; Fig 4E) despite declining betaine concentrations, suggesting consumption or export during osmotic adaptation. Alternatively, these betaine synthesis genes may be transcribed during desiccation but not translated or active until more energy becomes available upon rehydration. In hydrated controls, betaine catabolism transcripts (Transcriptome Module 1; Fig 4A) increased in abundance alongside steeper betaine depletion, consistent with its use as a carbon or nitrogen source during starvation. Supporting betaine catabolism during combined stress, transcripts coding for sarcosine oxidase, glycine hydroxymethyltransferase, and serine dehydratase (betaine-to-pyruvate pathway) were enriched in Transcriptome Module 1 and more abundant in controls (Fig. 4A). Finally, Targeted Metabolite Module 3 compounds (Fig. 5C) remained depleted in desiccated cells but increased upon rehydration in treatment samples, while also increasing throughout the experiment in starving hydrated control cells. These may represent critical compounds for active metabolism, including N-acetyl-alpha-D-galactosamine, vanillic acid, 3-hydroxyphenylacetic acid, riboflavin, pyridoxine, thymine, 2,4-dihydroxypyrimidine-5-carboxylic acid, and kynurenic acid.

We further analyzed 3,349 untargeted metabolite features from combined positive and negative ion modes (1,424 negative ion mode features and 1,925 positive ion mode features). Temporal analysis revealed that 133 features (4%) exhibited significant temporal trajectories across the experimental conditions (Supplementary Table 3). These features were clustered into three modules based on their combined treatment and control patterns, mirroring the approach used in the targeted metabolomics analysis. Untargeted Metabolite Module 1 (Supplementary Fig. 3A) exhibited a temporal profile like Targeted Metabolome Module 1 (Fig. 5A): features increased during desiccation, returned to baseline upon rehydration, and remained stable under control conditions. Untargeted Metabolite Module 2 is comprised of features that remained stable in the treatment condition across days 2, 8, and 14 but decreased in intensity upon rehydration (Supplementary Fig. 3B). These same features were progressively depleted in controls, consistent with their loss in metabolically active hydrated cells while persisting in metabolically arrested desiccated cells. Untargeted Metabolite Module 3 features increased under control conditions but remained low in treated cells on days 2, 8, and 14 (Supplementary Fig. 3C), then increased markedly upon rehydration—consistent with their production or accumulation in metabolically active hydrated cells.

Many untargeted features could not be assigned specific molecular formulas. Among those that were assignable, several showed consistent patterns across both targeted and untargeted analyses. For example, gluconic acid, uridine, and hypoxanthine exhibited similar temporal trajectories in both datasets (Fig 5A and Supplementary Fig. 3A). In contrast, several metabolites that showed strong temporal signals in the targeted analysis—with increased intensity upon desiccation and return to baseline upon rehydration (Fig. 5A)—exhibited more muted responses in the untargeted analysis (Untargeted Metabolite Module 2; Supplementary Fig. 3B), characterized by stable intensity during desiccation rather than increases, and reduced intensity in hydrated cells. These metabolites included adenosine, deoxyuridine, nicotinamide, adenine, xanthine, and methylthioadenosine. The differences between analytical approaches likely reflect methodological factors: targeted analyses use compound-specific optimization and isotope-labeled internal standards for each metabolite, providing greater sensitivity and accuracy for known compounds, while untargeted approaches employ generic parameters optimized for detecting the broadest possible range of features, often at the expense of sensitivity for specific metabolites. The targeted data are therefore more reliable for quantifying these specific compounds, while the untargeted data provide complementary discovery of unanticipated metabolites. Importantly, despite quantitative differences, the directional trends—depletion in active cells and persistence in dry cells—were generally consistent between approaches.

## CONCLUSION

Dormancy is defined as a reversible state of low to no metabolic activity (68). Our results demonstrate that desiccation induces a dormant state in *Arthrobacter* sp. AZCC_0090, characterized by compositionally stable mRNA and intracellular metabolite profiles in the desiccated state (days 8-14), despite dramatic restructuring during the transitions into and out of desiccation. Figure 6 synthesizes these findings as a conceptual diagram of molecular responses across the desiccation-rehydration continuum, illustrating how physiological processes sequentially activate during drying, stabilize in dormancy, and reactivate upon rehydration.

**Figure 6:**
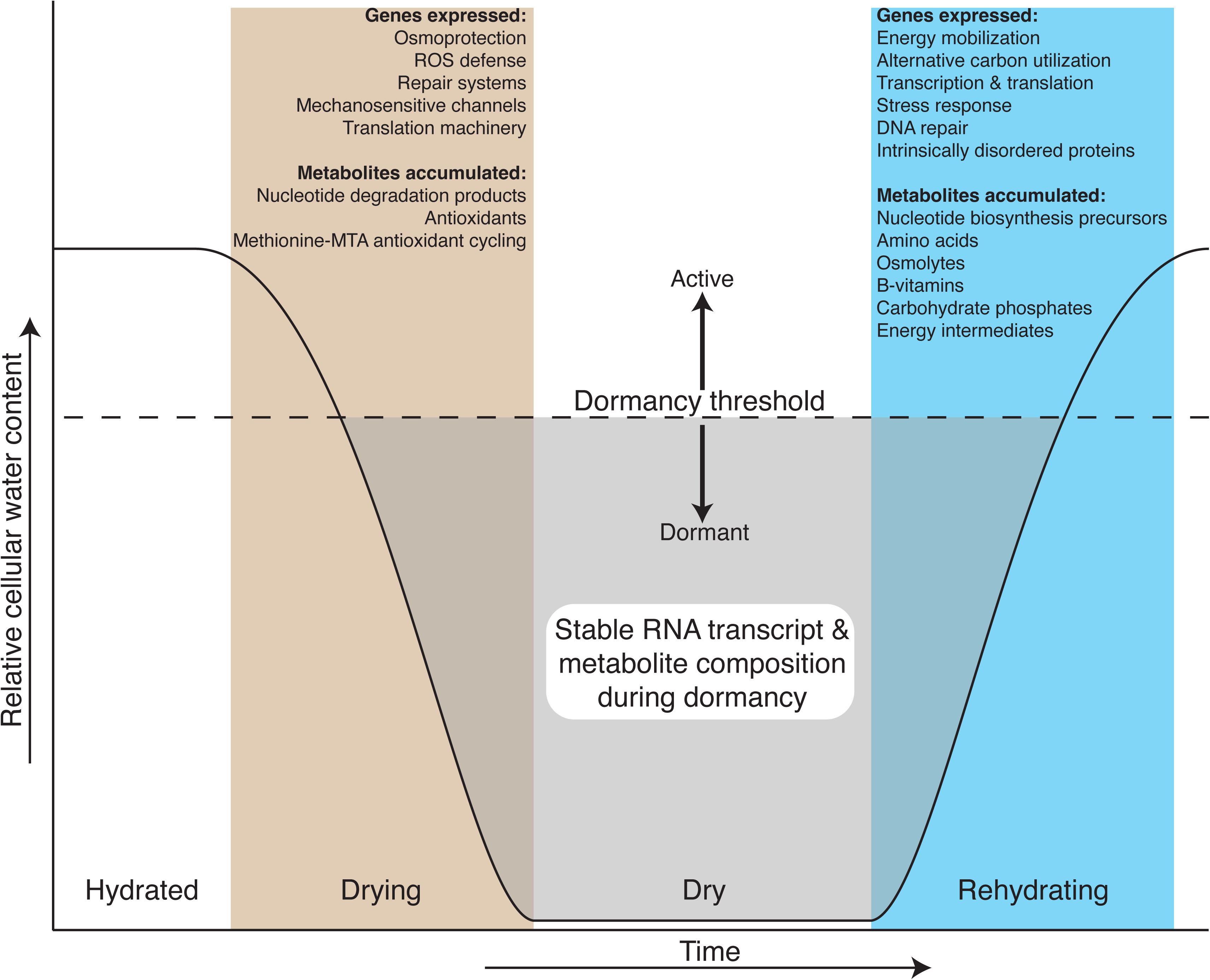
Conceptual diagram showing how *Arthrobacter* molecular processes respond across the desiccation-rehydration continuum. As cellular water content decreases, cells sequentially activate protective mechanisms and accumulate stress-related metabolites. Below the dormancy threshold (dashed line and shaded area), RNA transcript and metabolite composition stabilizes, indicating metabolic quiescence. During rehydration, cells suppress desiccation responses and activate energy mobilization, biosynthesis, and growth machinery.

Water vapor was sufficient to resuscitate dormant cells, inducing shifts in both mRNA and metabolites that prime cells for regrowth. We identified the accumulation of ribonucleosides and nucleobases during desiccation that persisted when mRNA profiles were stable. The source of these nucleotide degradation products during a period of apparent transcriptional stability remains unclear, but may represent degradation of damaged RNAs, turnover of non-coding RNAs not captured in our analysis, or accumulation of RNA turnover products generated during the drying transition. More focused experiments measuring RNA turnover rates directly are needed to resolve this apparent paradox.

The signatures we observed in desiccation-induced dormancy differ markedly from other dormancy states such as endospores. While most RNAs in dormant endospores are unstable and degrade to serve as nucleotide reservoirs for germination (44, 69), *Arthrobacter* maintained stable RNA profiles throughout desiccation. This pattern more closely resembles RNA dynamics in the non-spore forming actinobacterium, *Curtobacterium*, and desert communities where RNA profiles remained stable over extended dry periods (43, 70, 71).

Our findings challenge fundamental assumptions underlying RNA-based assessments of microbial activity in environments that experience desiccation. Microbial ecologists commonly use RNA presence as a proxy for microbial activity, based on the rationale that RNA is labile and would degrade in dead or inactive cells (48, 72). While RNA turnover is rapid in growing cells and partially applies to dormant endospores (44, 69), this assumption has not been systematically tested for desiccated non-spore forming bacteria. Methods applying rRNA:rDNA gene abundance ratios (51, 73) and metatranscriptomic analyses (71, 74, 75) operate under the assumption that extracted RNA reflects metabolically active populations. However, our data demonstrate that: 1) RNA is extractable and sufficiently intact for quantification and sequencing from desiccated cells, 2) some of these RNAs remain abundant during desiccation-induced dormancy, and 3) proportional transcript changes occur primarily during transitions into or out of desiccation, but not during the dormant desiccated state.

Transcriptomes from desiccated cells may reflect the transcriptional state when cells entered dormancy, rather than ongoing transcription. Our results support this interpretation. The stable mRNA composition we observed from days 8 to 14 indicates transcriptional stasis, not active transcription in the dormant state. However, this presents a complex interpretational challenge: RNA extracted from environmental samples experiencing desiccation cannot distinguish between metabolically active cells and dormant cells retaining transcripts from their pre-dormancy state. This issue is particularly acute in soil studies, where extraction protocols that process 1-2 g soil inevitably capture microenvironments with varying hydration states, yielding RNA from mixed populations of dormant and active cells that cannot be easily disentangled using current methods. Consistent with this, recent community-level studies show transcriptional responses follow rewetting (36, 71, 76), suggesting the transcriptional signatured detected in field samples may represent a mixture of dormancy-associated transcripts and active transcription during a hydration event. The ability to extract compositionally stable RNAs from desiccated cells therefore suggests that RNA-based activity assessments may overestimate *in* situ microbial activity in dryland soils by conflating dormant cells retaining stable transcripts with metabolically active populations.

There are several important limitations of this work. First, we examined only one strain of *Arthrobacter*, a genus known for its desiccation tolerance. Thus, it is unclear whether RNA stability is a specific feature of *Arthrobacter*’s desiccation tolerance mechanisms or represents a more general response to desiccation stress. However, the similar findings in *Curtobacterium* isolates (70) and in complex microbial communities (36, 71) suggest this phenomenon may be widespread among desiccation-tolerant bacteria. Moreover, the stability of transcript and metabolic composition from days 8-14 suggests cells reach a functionally dormant state by day 8. Yet, the point at which water loss induces dormancy remains to be fully characterized. Future studies employing direct measurement of intracellular water content and high-resolution temporal sampling during early desiccation will be needed to establish these relationships. Additionally, these experiments were performed under controlled laboratory conditions, and their relevance field conditions remains yet to be determined. Unfortunately, methods for studying RNA stability in soil microbial communities are currently lacking. Finally, while we demonstrated water vapor was sufficient for resuscitation, the mechanisms by which desiccated cells sense and respond to water vapor remain unknown. Some metabolites accumulated during desiccation may exhibit humectant-like properties, but the biophysical and molecular basis of vapor-phase rehydration requires further investigation.

The persistence of RNA in desiccated cells has implications extending far beyond arid soil microbiology, with relevance for understanding microbial survival and dispersal in extreme environments, astrobiology applications, pathogen dispersal across climate regimes, and the function of airborne microbiomes. The ability of desiccated cells to maintain RNA while remaining dormant until water vapor triggers resuscitation suggests environmental microbes may be more resilient than previously recognized, potentially reshaping our understanding of microbial biogeography and ecosystem function in water-limited systems on Earth and beyond.

## METHODS

### Strain source and propagation

*Arthrobacter sp.* strain AZCC_0090 was isolated as described elsewhere (52) and its genome sequence is available (53). All growth experiments were conducted on Yeast Mannitol media (YM), consisting of (per liter) 1.0 g yeast extract, 10.0 g mannitol, 0.5 g dipotassium phosphate, 0.2 g magnesium sulfate, 0.1 g sodium chloride, and 1.0 g calcium carbonate. Solid media was prepared with 2% Noble agar. Cell washes and resuspensions were conducted in YM salts (per liter): 0.5 dipotassium phosphate, 0.2 g magnesium sulfate, and 0.1 g sodium chloride. All incubations were at 25°C.

### Cell preparation for desiccation

We pelleted a 1.0 L overnight (∼18 h) culture of AZCC_0090 (27,500 rpm for 30 minutes) and washed once with YM salts before resuspending in 1.0 L YM salts. We incubated resuspended cells overnight at 25°C with shaking. The following day, 10 ml aliquots of the cell suspension (∼5-6 1110^8^ cell ml^-1^) were vacuum-filtered through individual 0.2 µm pore-size hydrophilic non-hygroscopic polycarbonate membrane filters (Millipore Sigma Polycarbonate Membrane Filter GTTP02500; 25 mm diameter) to collect cells. Each filter was placed into a vented petri dish. For “washed” control samples, cells were filtered as described above but processed immediately for viability assays and total and ribosomal RNA quantification without incubation in the desiccation chamber.

### Humidity chamber construction and humidity control

Humidity chambers were constructed from plastic storage containers. Several 1/4” diameter ports were drilled in each container to accommodate humidity sensor wires and to suspend a small fan (SEPA 12V 0.03A). Access ports were sealed with tape or silicone. The relative humidity was controlled using a series of supersaturated salt solutions as described previously (77). The following saturated salts were prepared with Nanopure water (nH_2_O): nH_2_O (no salt added), KCl, NaCl, NaBr, K_2_CO_3_, MgCl_2_, and CH_3_CO_2_K. The salts were considered saturated by the appearance of precipitated salt at 25°C. Saturated salt solutions or water were contained in an open beaker sitting underneath a fan blowing toward the surface of the solution. The fans were wired to a power supply (TekPower TP3005T) set to 12V 0.08A. At each change in RH, the humidity chamber was opened and the beaker containing the water or saturated salt was exchanged and the humidity chamber was closed. Temperature and relative humidity were continuously monitored with a Sensirion SEK-SHT35 digital humidity sensor placed inside an empty, closed vented petri dish. The Sensirion sensor was connected to a laptop via a Sensiron SEK-SensorBridge running the Sensiron Control Center. Chambers were maintained at constant temperature (25°C) throughout the experiment to prevent temperature fluctuations that could cause condensation. Supplementary Fig 4 shows the real time temperature and RH data during the rehydration phase. Filters were visually inspected during sampling and showed no evidence of liquid water accumulation.

### Experimental Design

We randomized filtered cells to treatment or control humidity chambers.

The RH in the control chamber was maintained at 99-100% for the duration of the experiment. The RH in the treatment chamber was slowly dehydrated from 100% RH to ∼25% RH over the course of two weeks (48 h at each humidity level). After 14 days the treatment chamber was rehydrated with water vapor for 48 h. Replicate filters were selected randomly from the treatment and control chambers for cultivability assays, metabolomics, or transcriptomics at days 2, 8, 14, and 16 after the start of the experiment. The chambers were only opened to change the saturated salt solutions or for sampling.

### Cultivability and spore formation assays

Cells were resuspended from filters in YM salts by vortexing for 10 minutes at maximum speed. 200 µl of resuspended cells were fixed with 5% formaldehyde (vol/vol) and stained with SYBR green I (1:40 dilution of commercial stock in Tris-EDTA) for 24 hours before counting with an EMD-Millipore Guava Technologies flow cytometer (Millipore, Billerica, MA, USA), as described previously (52). Culturable cells were enumerated by plate counts by serially diluting resuspended cells in YM salts and plating on solid YM. We defined percent cultivability as the percent of resuspended cells enumerated with flow cytometry that formed colonies on solid media. We used a 1 h treatment with 70% ethanol to determine if *Arthrobacter* produced ethanol-resistant spores. In brief, filters containing dried cells were floated on 70% ethanol for 1 h before spotting on solid YM and incubating. No growth was observed after ethanol treatment.

### RNA extraction, sequencing and mapping

Replicate filters were placed into microcentrifuge tubes containing glass disruptor beads, and frozen at -80°C until extraction. RNA was extracted using a Qiagen RNEasy mini kit, per the manufacturer’s instructions. DNA was removed with Ambion TURBO DNase per the manufacturer’s instructions. RNA was quantified with a Qbit using hs or br kits, as appropriate based on RNA concentration. Samples were sequenced at SeqCenter, LLC (Pittsburgh, PA), using their standard protocols. In brief, RNAs were DNAse treated a second time with Invitrogen DNAse (RNAse free). Library preparation was performed using Illumina’s Stranded Total RNA Prep Ligation with Ribo-Zero Plus kit and 10bp IDT for Illumina indices. Samples were sequenced on a NextSeq2000 giving 2x51bp reads.

Demultiplexing, quality control, and adapter trimming was performed with BCL Convert v3.9.3. The quality of generated raw reads was assessed by FastQC v0.73 via Galaxy (galaxy0) (78, 79). The reads were mapped to *Arthrobacter* sp. AZCC_0090 genome using Bowtie2 v2.4.1 (80) with default parameters in paired-end mode. Counts were generated with the featureCounts function in the Rsubread v2.12.3 R Bioconductor package (81).

### rRNA qPCR

RNA extracts were used as template for RT-qPCR using a SuperScript™ III One-Step RT-PCR System with Platinum™ *Taq* DNA Polymerase kit, per the manufacturer’s instructions. 16S rRNA RT-qPCR was performed using 515F 5′-GTGCCA GCMGCCGCGGTAA-3′ and 806R 5′-GGACTACHVGGGTWTCTAAT-3′ 16S rRNA primers (82). 23s rRNA RT-qPCR was performed using 111F 5’-ATGTCCGAATGGGGAAACCC-3’ and 557R 5’-CACGGTACTGGTCCGCTATC-3’ 23S rRNA primers that were designed for this study from the *Arthrobacter* sp. AZCC_0090 genome sequence (53). Cycle conditions for both 16S and 23S rRNAs: 48°C 30 minutes, 95°C for 10 minutes, followed by 40 cycles of 95°C 15 seconds, and 60°C for 1 minute. All qPCRs were run in triplicate and quantified against a standard curve prepared with *Arthrobacter* AZCC_0090 gDNA.

### Temporal analysis of RNA abundances

Differential expression analysis was performed using ImpulseDE2 (83) to identify genes with distinct expression across the experimental conditions. ImpulseDE2 was selected for its ability to model complete temporal trajectories and detect impulse-like expression patterns where genes temporarily change expression before returning to baseline levels (83). The analysis was conducted in case-control mode to compare treatment (case) and control conditions across all time points simultaneously, using count data for genes that had at least 10 reads in three or more samples (3709 of 4653 genes). Genes with adjusted p-values ≤ 0.05 were considered significantly differentially expressed.

### Clustering for temporal transcriptome dynamics

We used DESeq2 size factor normalized expression values for significantly differentially expressed genes from the ImpulseDE2 analysis for clustering analysis. Size factor-normalized expression values were standardized using Z-score normalization to ensure gene transcripts were grouped by expression patterns rather than magnitude. Mean expression values were calculated for each gene at each time point within treatment and control conditions. This double normalization precludes statistical comparisons between time points. To determine the optimal number of gene expression clusters, we used hierarchical clustering using correlation-based distances (1 - Pearson correlation) and Ward’s linkage. The number of clusters was determined using the elbow method applied to within-cluster sum of squares, with a minimum threshold of 6 modules to ensure adequate resolution of expression patterns.

### Metabolite extraction

Replicate filters were frozen at -80°C until metabolite extraction. Intracellular metabolites were extracted from filtered cells by direct sonication in methanol. Briefly, we added 1 mL of cold methanol to tubes containing filtered cells on ice and sonicated with a probe-style sonicator (Fisher Scientific Model FB120 with CL-18 probe) for 5 minutes at an amplitude of 60%. Filter debris was separated from supernatant by centrifugation. The supernatant was transferred to a new tube and dried to completeness in an Eppendorf Vaccufuge plus at 45°C for 40 min. We extracted metabolites from several filters without cells and methanol only as negative controls.

### Metabolomics

Analyses were performed at the Joint Genome Institute using standard procedures. Dried extracts were resuspended in methanol containing internal standards (^13^C-^15^N labeled amino acids and 2-amino-3-bromo-5-methylbenzoic acid), filtered, and analyzed by LC-MS using an Agilent 1290 Infinity LC system (Agilent, Santa Clara, CA) coupled to a Thermo QExactive HF orbitrap mass spectrometer (Thermo Scientific, San Jose, CA). Polar metabolites were separated using HILIC chromatography on a Poroshell 120 HILIC-Z column with a water-acetonitrile gradient. Full MS spectra (m/z 70-1050) and MS/MS fragmentation data were acquired in both positive and negative ion modes. Sample injection order was randomized with methanol blanks between samples. Complete analytical conditions are provided in Supplementary Methods.

For targeted metabolite identification, experimental mass spectra were compared to compound standards using custom Python code (84). Spectral features were assigned confidence scores (0-3) according to Metabolomics Standards Initiative levels (85), with "level 1" identifications requiring mass accuracy and retention time matches to pure standards, and highest confidence identifications also requiring matching MS/MS fragmentation patterns. Complete analytical and identification parameters are provided in Supplementary Methods.

For untargeted features, experimental mass spectra of unassigned features with MS/MS fragmentation data were compared against the Global Natural Products Social Molecular Networking (GNPS) (86) that performs molecular networking and grouping partially and fully identified metabolites based on their MS/MS fragmentation tree, leading to the discovery of related metabolites/features based on common/similar functional groups; and SIRIUS 4 (87) which uses isotope patterns in MS spectra and fragmentation profiles in MS/MS spectra to further assist in molecular formula assignment and potential structural elucidation. CSI:FingerID was used for molecular structure annotation through database searches and CANOPUS for de novo compound class prediction. This integrated workflow supports comprehensive identification of both known and novel compounds from tandem mass spectrometry data in complex matrices.

### Volume adjustment and Ion mode selection

We experienced sample volume loss during metabolite extraction for nine samples before drying: T0C rep 4 (100 μL collected), T0C rep 1 (200 μL), T0E rep 5 (200 μL), T0E rep 4 (250 μL), T0E rep 3 (300 μL), T1C rep 3 (450 μL), T1E rep 3 (450 μL), T1E rep 1 (300 μL), and T2E rep 1 (400 μL). To account for this volume loss, we adjusted metabolite peak heights by multiplying by the ratio of expected volume (500 μL) to actual volume collected: Adjusted peak height = Original peak height × (500/Actual volume collected). For targeted metabolites identified in both positive and negative ion modes, we selected the polarity with higher mean peak heights for subsequent analyses. For untargeted features, duplicate metabolites in the negative and positive ion mode were identified. Features with identical predicted molecular formulas and retention times within 5 seconds across positive and negative ion modes were considered duplicates. For each duplicate pair, the metabolite measured in the ion mode with higher mean intensity across all samples was retained. This deduplication process removed 32 duplicated features. For both targeted and untargeted metabolomics, a small value was added to missing and zero values (10% of the smallest positive peak area) to facilitate Log transformation in downstream analyses.

### Temporal analysis of metabolites

ImpulseDE2 was designed for RNA-seq count data analysis but was adapted for metabolomics peak intensity data. To do this, we Log2-transformed the peak heights, multiplied them by 100, and rounded to create pseudo-count values. Results were extracted and clustered using the same framework as transcriptomics analysis described above. Significantly different metabolites were defined as those with Impulse2DE adjusted p-values ≤ 0.05.

### MDS plots and statistics of transcriptome and metabolome data

We used multidimensional scaling (MDS) plots and permutational multivariate analysis of variance (PERMANOVA) to compare transcriptome and metabolome profiles over time across conditions. Raw transcriptome count data were processed using DESeq2 (88) with variance stabilizing transformation for visualization and multivariate analyses. We Log10-transformed the metabolome data. For both datasets, multidimensional scaling was used to visualize sample relationships based on Euclidean distances. PERMANOVAs were implemented through the adonis2 function in the vegan R package (89), employing a factorial design (∼day + treatment + day:treatment) with 999 permutations. To resolve temporal dynamics within each condition, we conducted separate pairwise PERMANOVA comparisons between time point combinations for control and treatment groups independently.

## DATA AVAILIBILITY

Processed RNA-seq data generated in this study have been deposited in the NCBI Gene Expression Omnibus (GEO) under accession number GSE309279. Individual samples are found in accessions GSM9264299-GSM9264323. Raw metabolome data is available in the massIVE repository at the Global Natural Product Social Molecular Networking (GNPS) site under the accession MSV00009260 (free registration required). Code used in analyses and metabolite sample metadata are available at 10.6084/m9.figshare.30370339.

## ACKNOWLEDGEMENTS

The authors would like to thank Bradley Schlottman, and Caitlin Tribelhorn for assistance with equipment and experiments. Funding support is from the National Science Foundation in a Graduate Research Fellowship Grant award to IAV DGE-2137419 and IOS-2141605 to PC. PC, LM and PM were supported by the University of Arizona Research, Innovation & Impact (RII) and Technology Research Initiative Fund/Water, Environmental, and Energy Solutions. The work (proposal:https://doi.org/10.46936/10.25585/60001177) conducted by the U.S. Department of Energy Joint Genome Institute (https://ror.org/04xm1d337), a DOE Office of Science User Facility, is supported by the Office of Science of the U.S. Department of Energy operated under Contract No. DE-AC02-05CH11231.

