## Supplemental Figures for "Transcriptional and metabolic stasis define desiccation-induced dormancy in the soil bacterium *Arthrobacter* sp. AZCC_0090 until water vapor initiates resuscitation"

**Supplementary Table 1:** Full Table of genes and their impulseDE2 output and module clustering results

**Supplementary Table 2:** Full Table of targeted metabolites and their impulseDE2 output and module clustering results

**Supplementary Table 3:** Full Table of untargeted features and their impulseDE2 output and module clustering results

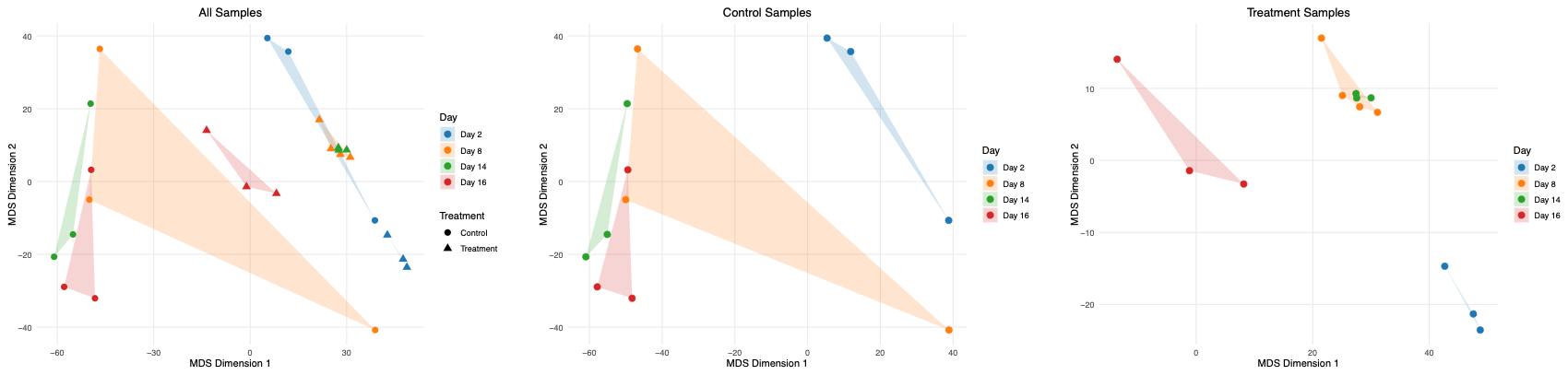

**Supplemental Figure 1: Multidimensional scaling analysis of transcriptome data. MDS plots based on variance-stabilized transformed RNA-seq data show sample clustering patterns across the 16-day desiccation experiment. (Left) All samples colored by timepoint (Days 2, 8, 14, 16) and shaped by treatment (circles = control, triangles = desiccation treatment). (Center) Control samples only. (Right) Treatment samples only. Points represent individual biological replicates, and shaded hulls around timepoint groups.**

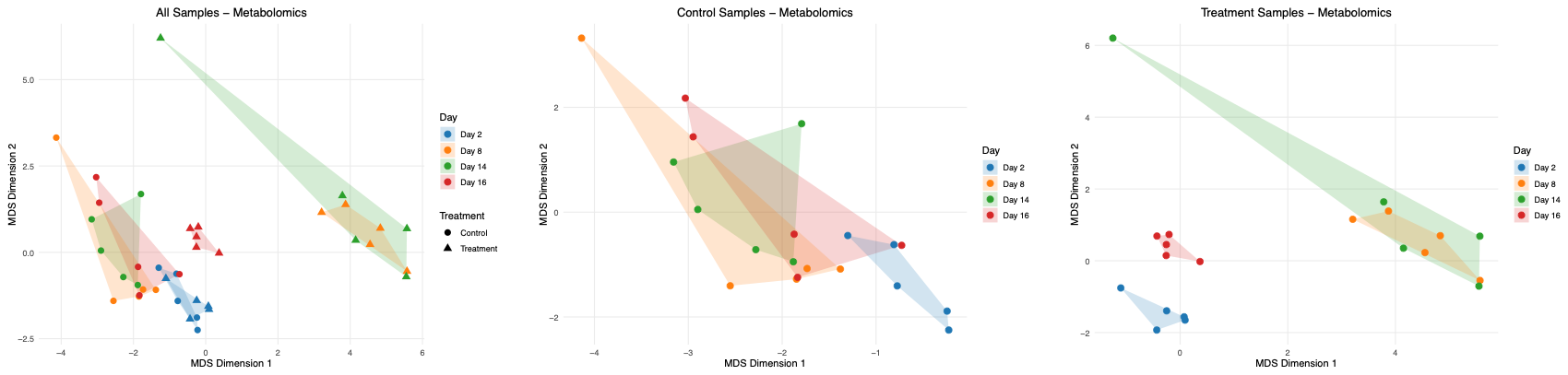

Supplemental Figure 2: **Multidimensional scaling analysis of targeted metabolome data reveals temporal and treatment-dependent patterns in gene expression profiles.** MDS plots based on log10-transfoprmed peak heights show sample clustering patterns across the 16-day desiccation experiment. (Left) All samples colored by timepoint (Days 2, 8, 14, 16) and shaped by treatment (circles = control, triangles = desiccation treatment). (Center) Control samples only. (Right) Treatment- samples only. Points represent individual biological replicates, and shaded hulls around timepoint groups.

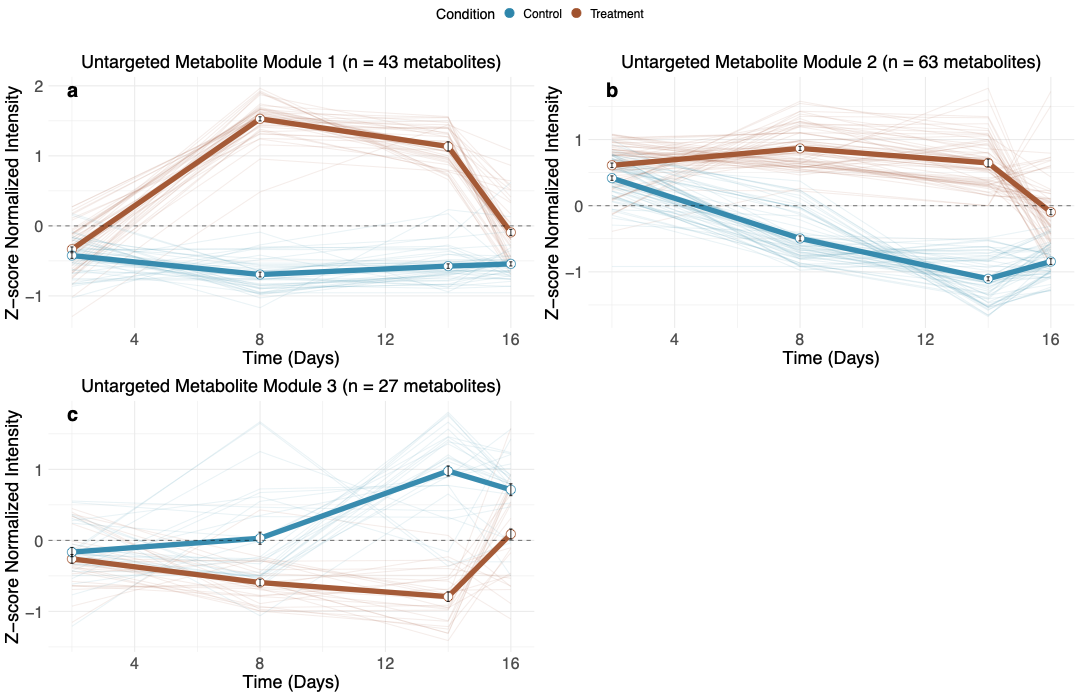

Supplementary Figure 3: Temporal expression modules of differentially abundant untargeted features**.** Three untargeted modules (a-c) were identified through correlation-based hierarchical clustering of untargeted features with significantly distinct temporal responses across treatment and control conditions (modified ImpulseDE2, FDR-corrected p ≤ 0.05). Each panel displays mean z-score normalized untargeted feature peak height profiles (±SEM) for control (bold blue) and treatment (bold brown) conditions, with individual untargeted feature profiles shown as thin lines. Sample sizes (n) indicate the number of untargeted features within each module. See supplementary table 3 for full ImpluseDE2 results and module assignments.

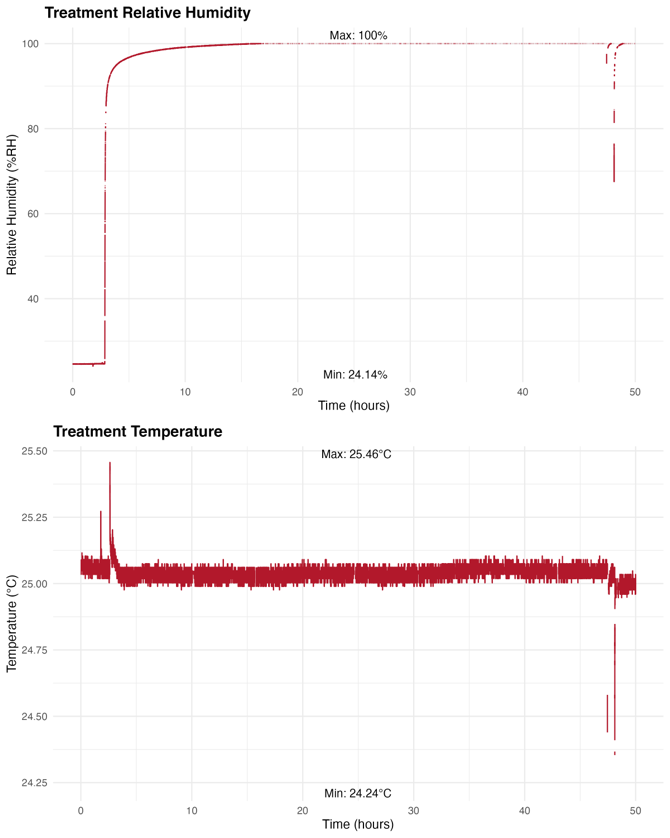

**Supplementary Figure 4: Temperature and relative humidity monitoring during rehydration.** Relative humidity (top) increased from 24.14% to 100% within approximately 15 hours following replacement of saturated salt solution with open water at time zero. Temperature (bottom) remained stable throughout the 48-hour period with minimal variation mostly explained by opening the chamber (minimum = 24.24°C, maximum = 25.46°C). Brief transient spikes in both parameters correspond to chamber opening events for sample manipulation. Stable temperature conditions indicate minimal risk of sustained condensation during rehydration. Note, the x-axis time scale is relative to the extracted data shown here, not total eclipsed time.

**Supplemental Methods:**

Metabolomics full details: Metabolomic analysis and compound identification were performed at the Joint Genome Institute, using their standard procedures. Briefly, dried extracts were resuspended in 170 µL 100% methanol containing a mixture of internal standards (15 µM, mix of 13C-15N labeled amino acids, #767964; 1 µg/mL, 2-Amino-3-bromo-5-methylbenzoic acid, #R435902).  Extracts were centrifuge-filtered (2.5 min at 2,500 rpm) through a 0.22 µm hydrophilic-PVDF membrane (UFC40GV0S, Millipore) and transferred to a glass vial.  Liquid chromatography mass spectrometry (LC-MS) analysis was performed using an Agilent 1290 Infinity LC system (Agilent, Santa Clara, CA) coupled to a Thermo QExactive HF orbitrap mass spectrometer (Thermo Scientific, San Jose, CA).  To detect polar metabolites using normal phase chromatography, 2 µL of each sample extract was injected onto a HILICZ column (Agilent InfinityLab Poroshell 120 HILIC-Z, #673775-924, 150x2.1mm, 2.7 µm) warmed to 40°C with a flow rate of 0.45 mL/min.  For metabolite separation, the column was equilibrated with 100% buffer B (95:5 acetonitrile:water w/ 5 mM ammonium acetate) for 1 minute, followed by a linear gradient diluting buffer B down to 89% with buffer A (100% water w/ 5 mM ammonium acetate and 5 µM methylene-di-phosphonic acid) over 10 minutes, then down to 70% B over 4.75 minutes, then down to 20% B over 0.5 minutes, and then isocratically held at 20% B for 2.25 minutes. Full MS spectra were acquired between m/z 70-1050 at 60,000 resolution, in centroid format in both positive and negative ion mode, with MS/MS fragmentation data acquired using average stepped collision energies of 10, 20, and 40 eV, as well as 20, 50 and 60 eV, at 15,000 resolution.  Mass spectrometer source settings included a sheath gas flow rate of 55 (au), auxiliary gas flow of 20 (au), sweep gas flow of 2 (au), spray voltage of 3 kV and capillary temperature of 400 ºC.  Sample injection order was randomized, with an injection blank of 100% methanol run between each sample.

To identify metabolites, experimental mass spectra including m/z, retention time and MS/MS fragmentation pattern were compared to that of compound standards run using the same LC-MS/MS methods.  Analysis was performed using custom Python code (84). Spectral features (unique m/z coupled with retention time, RT) were assigned a score from 0 to 3, representing the level of confidence in compound identification and then rated according to the Metabolomics Standards Initiative (MSI) confidence levels (85).  For compounds detected at m/z </= 5 ppm or 0.001 Da from theoretical as well as RT </= 0.5 min compared to a pure standard run using the same LC-MS method, a positive “level 1” identification was given.  A compound with the highest level of positive identification (score of 3), “exceeding level 1”, also had matching MS/MS fragmentation spectra in comparison to either an outside database or internal database generated from standards run and collected on a Q Exactive Orbitrap HF MS.  An identification was invalidated if an MS/MS mismatch was observed.
